# Allele Mining in Diverse Accessions of *Urochloa* and *Megathyrsus* spp. Tropical Grasses to Improve Forage Quality and Reduce Environmental Impact

**DOI:** 10.1101/2020.12.09.418087

**Authors:** SJ Hanley, TK Pellny, JJ de Vega, V Castiblanco, J Arango, PJ Eastmond, JS Heslop-Harrison, RAC Mitchell

**Affiliations:** Rothamsted Research, Harpenden, Hertfordshire AL5 2JQ, UK; Earlham Institute, Norwich Research Park, NR4 7UZ, UK; International Center for Tropical Agriculture (CIAT), 6713 Cali, Colombia; Department of Genetics & Genome Biology, University of Leicester, Leicester LE1 7RH, UK

## Abstract

The C4 *Urochloa* spp (syn. *Brachiaria*) and *Megathyrsus maximus* (syn. *Panicum maximum*) are used as pasture for cattle across vast areas in tropical agriculture systems in Africa and South America. A key target for variety improvement is forage quality: enhanced digestibility could decrease amount of land required per unit production and enhanced lipid content could decrease methane emissions from cattle. For these traits, loss-of-function (LOF) alleles in known gene targets are predicted to improve them, making a reverse genetics approach of allele mining feasible. We studied allelic diversity of 20 target genes (11 for digestibility, 9 for lipid content) in 104 accessions selected to represent genetic diversity and ploidy levels of *U. brizan*tha, *U. decumbens, U. humidicola, U. ruziziensis* and *M. maximum*. We used RNAseq and then bait-capture DNA-seq to improve gene models in a *U. ruziziensis* reference genome to assign polymorphisms with high confidence. We found 953 non-synonymous polymorphisms across all genes and accessions; within these, we identified 7 putative LOF alleles with high confidence, including ones in the non-redundant SDP1 and BAHD01 genes present in diploid and tetraploid accessions. These LOF alleles could respectively confer increased lipid content and digestibility if incorporated into a breeding programme.

**Highlight:** We found gene variants in a collection of tropical grasses that could help reduce environmental impact of cattle production.

## Introduction

The environmental impact of cattle production could be decreased by reducing the amount of land required *e.g* land sparing) and amount of methane (CH_4_) emitted per unit production (*i.e* emission intensity). This could be achieved by genetic improvement of pasture grass on which they feed: an increase in digestibility and energy content would allow the same production to be achieved on a smaller land area and an increase in lipid content in vegetative matter would decrease CH_4_ emitted per unit production, provided that these two traits could be improved without negative side effects, such as reduced growth or susceptibility to biotic or abiotic stresses.

Breeding of commercial tropical forage grass varieties in diploid and polyploid species and interspecific hybrids of *Urochloa* has been achieved by recurrent selection over many years, identifying superior-performing populations for key traits such as biomass production in different environments, resistance to pests and digestibility (Worthington and Miles, 2014). These targets increase efficiency of forage grass, such that less land is required for production. Increasingly, environmental targets such as decreased nitrogen losses (Nuñez *et al*., 2018; Villegas *et al*., 2020) and reduced methane emissions from grazing cattle (Gaviria-Uribe *et al.*, 2020) have become public breeding targets for improved pasture grasses. Continued improvement will be accelerated using genetic diversity that is available from accessions of the same genus available in genebank collections; the ploidy and relatedness of 280 of accessions from the International Center for Tropical Agriculture (CIAT) collection of *Urochloa* spp and genome composition of some of the polyploids (P. Tomaszewska, *pers. comm*.). The relation between these species has previously been studied using microsatellite markers (Triviño *et al*., 2017).

However, introduction of such material into current breeding programmes is a major undertaking requiring evidence of likely benefit; breeding of *Urochloa* tropical forage grasses is particularly complicated by obligate outcrossing in sexual accessions and occurrence of apomixis in half of the progeny (Worthington and Miles, 2014). For traits where there are known key genes and an understanding of how variants of these might affect the traits, an allele mining approach may be feasible where sequencing of target genes rather than phenotyping can be used to find potentially useful alleles. This reverse genetic approach can find useful loss-of-function variation that would not be found by phenotyping as its effect is hidden due to gene redundancy (Comai, 2005), particularly in polyploid or highly heterozygous material such as *Urochloa*, and provides the basis for perfect markers in crosses for following the alleles. Successful examples of allele mining for natural variation in known genes include studies on rice germplasm for starch synthesis genes (Butardo *et al*., 2017) and on Sorghum germplasm for a gene responsible for Al tolerance (Hufnagel *et al*., 2018).

Two traits where loss-of-function (LOF) alleles have been identified as beneficial are (1) digestibility where improvements have been gained by knock-out or knock-down of genes involved in cell wall synthesis in grasses and (2) lipid content of vegetative tissue where improvements could be gained by knock-out or knock down of genes involved in lipid metabolism. Increased lipid content of vegetative tissue of forage results in decreased CH_4_ emissions from cattle that feed on it, as well as benefiting meat and dairy fatty acid composition, as demonstrated by a transgenic approach in *Lolium perenne* (Winichayakul *et al*., 2020).

Here we compile a list of genes identified as targets from work in our labs or elsewhere, and find the orthologues in *U. ruziziensis* diploid reference species. We then conduct a comprehensive screening of alleles for these genes in 104 diverse accessions of *U. brizantha, U. decumbens, U. humidicola, U. ruziziensis* and *M. maximum* using RNAseq and bait capture genomic DNA sequencing.

## Materials and Methods

The sections below correspond to the steps shown in red in Figure 1.

**Figure 1.**
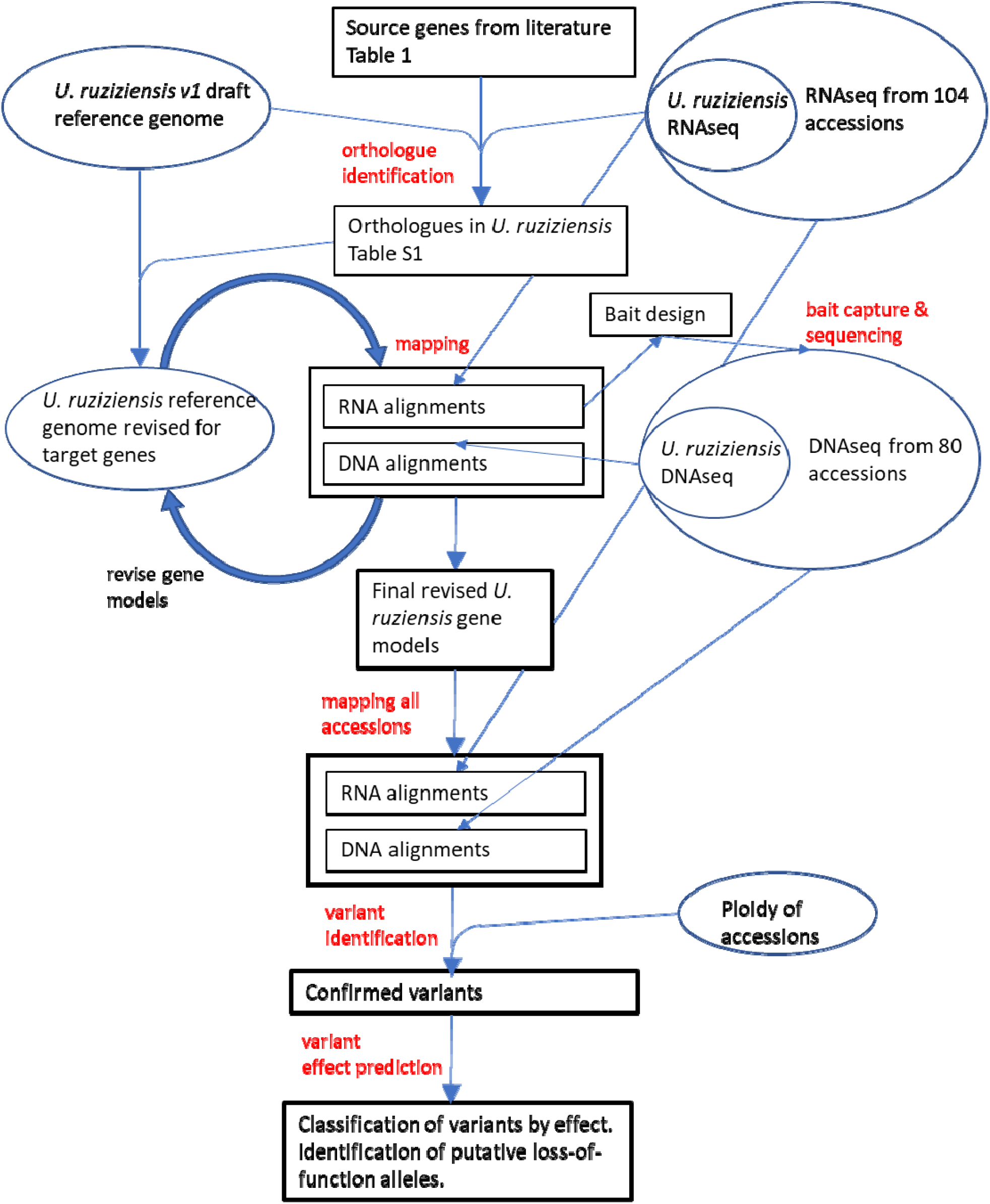
Workflow for analyses. Steps in red text are described in Methods section.

### Plant materials and RNA sequencing

We collected leaf samples of 104 accessions (Supplemental Table 1) from the field-grown genebank collection at CIAT which were immediately frozen in liquid nitrogen. Samples were ground to a fine powder in liquid nitrogen and subsequently lyophilised. Total RNA was extracted as described in (Pellny *et al*., 2012) with the difference that prior to DNAse treatment the pellets were dried in a rotary evaporator (Eppendorf) and stored/transported at room temperature. Illumina sequencing using standard RNA-seq library preparations with paired reads of length 150 bp was conducted by Novogene, HK. The raw reads were deposited in SRA under Bioproject PRJNA513453. We also collected leaf samples from an overlapping set of accessions for DNA extraction (Supplemental Table 1) and 80 of these were used for DNA sequencing described below.

### Orthologue Identification

We searched for *U. ruziziensis* orthologs of the target genes identified in other species listed in Table 1. Firstly, we identified the putative orthologues of the target genes in *Setaria viridis* (Setaria) v1.1 genome (Goodstein *et a*l., 2012) as the closest reliably annotated genome using BLASTN with target genes’ CDS as original queries and source genomes (i.e. Arabidopsis, maize, sorghum, Setaria or Brachypodium) of target genes for reciprocal BLASTN of hits. This identified 1-to-1 orthologues for all source genes except for Arabidopsis SDP1 and PXA1 genes where there were two putative paralogues each in Setaria. We repeated this process for the draft *U. ruziziensis* v1.0 annotated genome (Worthington *et al*., 2020) and found the same orthologous relationships as for Setaria except for one target gene CGI58 lipase, where an additional paralogue was found in *U. ruziziensis* v1.0. We compared the *U. ruziziensis* v1.0 gene models with the corresponding Setaria and source genes to judge whether they were correct; for 10 of 22 they were incomplete. We compiled a set of 22 genes using the 12 complete *U. ruziziensis* v1.0 genes and 10 Setaria genes for the others. We then mapped RNAseq reads from 11 *U. ruziziensis* accessions to this set. Two *U. ruziziensis* v1.0 genes with no equivalents in Setaria had almost zero mapped reads, and we removed these as likely pseudogenes, leaving a total of 20 *U. ruziziensis* target genes. Baits were designed to these 20 *U. ruziziensis* genomic regions taking account of the mapped RNAseq to customise baits for each species-ploidy group. Bait capture was performed on genomic DNA isolated from 80 accessions. Resulting IonTorrent sequencing, RNAseq reads and targeted Sanger sequencing for *U. ruziziensis* accessions were together used to check and refine gene models. We annotated the CDS by finding the longest ORF and comparing with that of orthologues. For 19 of these, we found complete coding sequences, but gene Ur.CGI58 lacks the first exon. We deposited final annotated sequences for all 20 *U. ruziziensis* genes in Genbank/EMBL accession numbers (MW323383-MW323402).

**Table 1.** Evidence from literature for selection of target genes to improve forage quality.

| target trait | gene | species | suppression mode | Effect on trait | pleiotropic effects | Ref. |
| --- | --- | --- | --- | --- | --- | --- |
| Digest-<br>ibility | 4CL /<br>Class I | *Sorghum*<br>*bicolor* | missense<br>mutant<br>bmr2 | 17% increased<br>saccharification |  | (Saballos *et al.*, 2008;<br>Sattler *et al.*,<br>2010) |
|  |  | *Saccharum*<br>*officinarum* | RNAi | 52%, 76 %<br>improved<br>saccharification<br>(field-grown) | 0%, 30 %<br>DM yield<br>penalty<br>(field-grown) | (Jung *et al.*,<br>2016) |
|  | BAHD01 | *Setaria viridis*,<br>*Saccharum*<br>*officinarum* | RNAi | 40-80% increased<br>saccharification | no growth<br>penalty in<br>GH | (de Souza *et al.*, 2018; de<br>Souza *et al.*,<br>2019) |
|  | BAHD05 | *Setaria viridis* | RNAi | 10-20% increased<br>saccharification | no growth<br>penalty in<br>GH | (Mota *et al.*,<br>2020) |
|  | COMT | *Zea mays* | bm3 LOF<br>mutant | used in<br>commercial<br>hybrids with<br>improved<br>digestibility for<br>cattle | some<br>reports<br>yield<br>penalty,<br>sometimes<br>no yield<br>penalty | (Sattler *et al.*,<br>2010; Vignols<br>*et al.*, 1995) |
|  |  | *Sorghum*<br>*bicolor* | bmr12 LOF<br>mutant | 30% increased<br>tract digestibility | 10% DM<br>yield<br>penalty | (Saballos *et al.*, 2008;<br>Sattler *et al.*,<br>2010) |
|  |  | *Panicum*<br>*virgatum* | RNAi | 30% increased<br>digestibility | no growth<br>penalty in<br>GH | (Fu *et al.*,<br>2011a) |
|  |  | *Saccharum*<br>*officinarum* | TALEN<br>induced LOF<br>mutations in<br>multiple<br>paralogs | 40 % improved<br>saccharification<br>(field-grown) | no DM<br>yield<br>penalty<br>(field-grown) | (Kannan *et al.*,<br>2018) |
|  | CAD /<br>Group I | *Brachypodium*<br>*distachyon* | missense<br>mutants | 40% increased<br>saccharification | no growth<br>penalty in<br>GH | (Bouvier<br>d'Yvoire *et al.*,<br>2013) |
|  |  | *Sorghum*<br>*bicolor* | missense<br>mutant<br>bmr6-3 | 20% increased<br>tract digestibility | 15% DM<br>yield<br>penalty | (Saballos *et al.*, 2009;<br>Sattler *et al.*,<br>2010) |
|  |  | *Zea mays* | bm1 |  | no yield<br>penalty | (Halpin *et al.*,<br>1998; Sattler<br>*et al.*, 2010) |
|  |  | *Panicum virgatum* | RNAi | 20% increased saccharification | normal GH growth | (Fu *et al.*, 2011b) |
|  | CCR | *Zea mays* | RNAi | 20% increased saccharification | increased growth GH | (Park *et al.*, 2012) |
|  | GT43A / IRX14 | *Brachypodium distachyon* | missense mutant? | 10% increased saccharification | no growth penalty in GH | (Whitehead *et al.*, 2018) |
|  | CCoAOMT | *Zea mays* | association with polymorphisms | correlation with fibre digestibility |  | (Brenner *et al.*, 2010) |
| lipid content | SDP1, SDP1-like | Arabidopsis | LOF mutant | increase in leaf triacylglycerol | poor seedling establishment in oilseeds | (Kelly *et al.*, 2013) |
|  |  | *Medicago truncatula* | VIGS | Increase in leaf lipid content | none | (Wijekoon *et al.*) |
|  | CGI-58 | Arabidopsis | LOF mutant | increase in leaf triacylglycerol | none | (James *et al.*, 2010) |
|  | PXA1, PXA1-like | Arabidopsis | LOF mutant | increase in leaf triacylglycerol | poor seedling establishment in oilseeds, starvation sensitive, jasmonate deficient | (Slocombe *et al.*, 2009) |
|  |  | *Medicago truncatula* | VIGS | Increase in leaf lipid content | none | (Wijekoon *et al.*) |
|  | TGD1 | Arabidopsis | leaky mutant | increase in leaf triacylglycerol | embryo defect, growth penalty | (Xu *et al.*, 2005) |
|  | TGD2 | Arabidopsis | leaky mutant | increase in leaf triacylglycerol | embryo defect, growth penalty | (Awai *et al.*, 2006) |
|  | TGD3 | Arabidopsis | leaky mutant | increase in leaf triacylglycerol | embryo defect, growth penalty | (Lu *et al.*, 2007) |

### Read Mapping

We did the mapping and variant calling on the Galaxy platform (Giardine *et al*., 2005). We first mapped RNAseq of 11 *U. ruziziensis* accessions to the *U. ruziziensis* v1.0 reference genome using BWA-MEM (Li and Durbin, 2010). Taking these alignments into Geneious, for each target gene, we combined mapped reads with set of all unmapped reads and conducted a *de novo* assembly. We compared resulting contigs with *U. ruziziensis* and Setaria viridis gene models and manually improved *U. ruziziensis* gene models. We substituted these gene models for the original versions as a first attempt at improving the reference and designed baits based on these. After we completed sequencing of bait capture DNA, we mapped DNA and RNA reads of *U. ruziziensis* accessions to the modified reference to iteratively improve it until all reads mapped satisfactorily. We substituted the final version of the *U. ruziziensis* gene models (18 out of 21 were changed from original version) into the *U. ruziziensis* genome annotation and mapped the RNA and DNA reads of all accessions using HiSAT2 (Kim *et al*., 2015) and the TMAP mapper within Torrent Suite 5.12.2 software (Thermofisher), respectively. For the latter, only reads greater than 100bp were used. We used the resulting BAM files (104 from RNAseq, 80 from DNAseq) to call variants and to manually inspect alignments on IGV (Robinson *et al*., 2011) for putative LOF alleles.

### Bait Capture

Coding sequences of the 20 genes of interest were targeted using myBaits Custom DNA-seq technology (Arbor Biosciences). A single bait set was designed to capture all genes in any of the species studied. To account for the likely diversity represented within and between species, consensus sequences were derived from the available RNAseq data for all genes in all individuals of each of the species. These were submitted to the design process performed by Arbor Biosciences which resulted in 20,346 baits of 70 nucleotide length with 3x tiling. Genomic DNA was extracted from frozen leaf tissue using a Plant DNeasy Kit (Qiagen) according to the supplied protocol. DNA quality was assessed by agarose gel electrophoresis and quantified using the Qubit dsDNA BR Assay Kit (Thermofisher). Whole genome libraries for use in bait capture were prepared using Ion Plus Fragment Library kits according to the manufacturer’s instructions with a target insert size of 400bp and unique Ion Xpress barcodes for each sample. Libraries were then amplified using the library kit PCR reagents to generate sufficient DNA for bait hybridisation. All libraries were quantified by qPCR using a Kapa Library Quantification Kit (Roche) and 16 equimolar pools made, each comprising five libraries. For each pool, bait capture was performed according to the manufacturer’s myBaits^®^ Manual v4. Libraries were then quantified by qPCR as before, pooled and sequenced across two runs on an Ion Torrent PGM sequencer, using Ion Hi-Q View OT2 reagents for 400bp templating and the Ion PGM Hi-Q View Sequencing Kit for 400 bp sequencing.

### Variant Calling

We used FreeBayes (Galaxy Version 1.0.2.29-3) to call variants, which is a haplotype-based variant calling program capable of dealing with polyploidy (Garrison and Marth, 2012). Both RNAseq and DNA capture BAM files (104 RNA, 80 DNA, overlap of accessions 74) were divided into 11 groups with the same species and ploidy (Table S2) using ploidy information from cytogenetics for the accessions (P. Tomaszewska, *pers. comm*.). BAM files from each group were submitted together to FreeBayes with the appropriate setting for ploidy, variant calling was limited to the target genes with default parameters for DNA reads; for RNA reads, the minimum fraction of observations supporting an allele (--min-alternate-fraction) was set to 0.05 to allow for low abundance due to nonsense-mediated decay of transcripts (Gutierrez *et al*., 1999) from LOF alleles. All other FreeBayes parameters were default. We retained variants with quality>=20 using SNPsift v4.0 (Cingolani *et al*., 2012a). Using custom Perl scripts, we compared polymorphisms from DNA and RNA VCF files produced by FreeBayes for the same group. Polymorphisms observed from the RNAseq were filtered out unless they were also observed in the corresponding DNAseq bait capture sequences for the same accession, or where this was not present, in another accession from the same species.

### Variant Effect Prediction

We identified effects on function of the putative polymorphisms with SNPeff v4.0 (Cingolani *et al*., 2012b), which uses the CDS annotation to predict effects on encoded proteins. We compiled information on all unique variants using custom Perl scripts to process VCF files (summarised in Table 2 and Figs 2, 3).

**Table 2.**
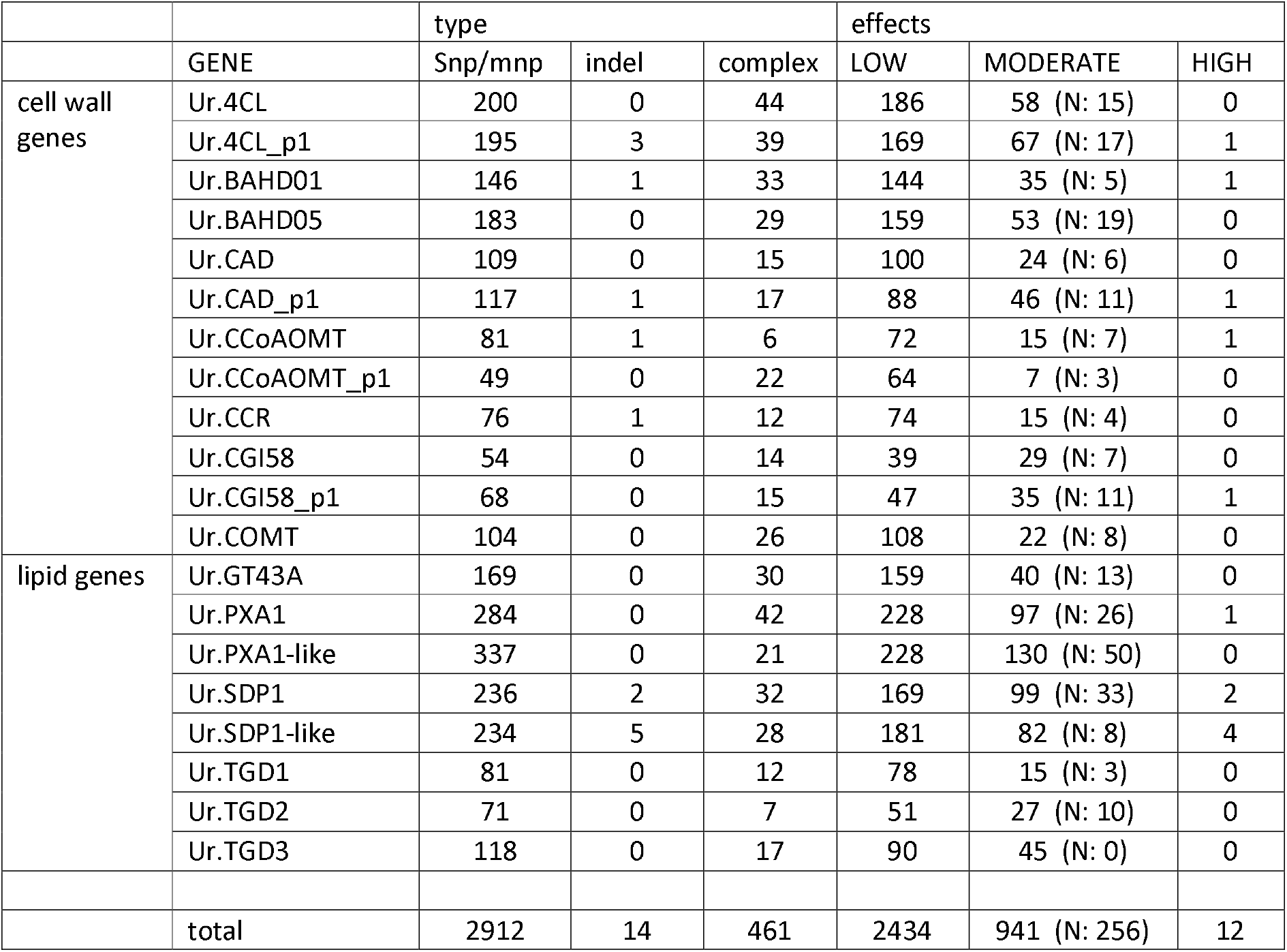
Total numbers of polymorphisms found in RNAseq and confirmed in bait capture DNA of 124 accessions for the 20 target *U. ruziziensis* genes. Genes are classified by type or predicted effect on protein. Type ‘snp/mnp’ includes SNPs and a small number of contiguous multiple nucleotide polymorphisms in the same haplotype; ‘complex’ denotes a mixture of SNPs and indels. ‘LOW’ effects are synonymous variants, ‘MODERATE’ are missense, in-frame indels, start or stop lost, and ‘HIGH’ are frameshift or stop gained, predicted to cause loss of function (LOF). From MODERATE missense variants, counts of those predicted as non-tolerated by SIFT web server are shown “(N:)”.

**Figure 2.**
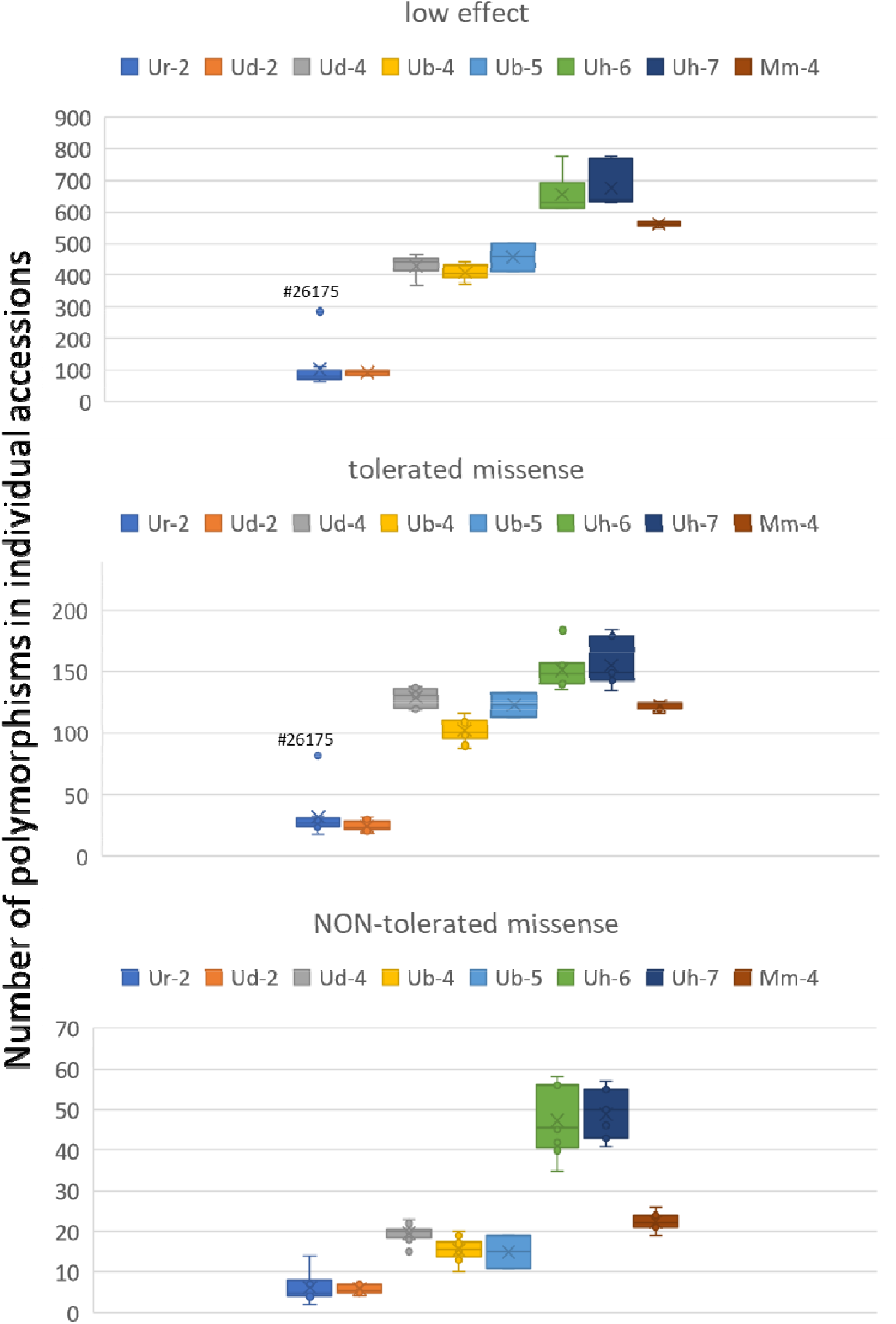
Number of variants in individual accessions grouped by species and ploidy. Only the 66 accessions with both RNA and DNA sequencing were used for this analysis. Ur-2 *U. ruziziensis* 2x, Ud-2 and Ud-4 *U. decumbens* 2x and 4x, Ub-5 and Ub-5 *U. brizantha* 4x and 5x, Uh-6 and Uh-7 *U. humidicola* 6x and 7x, Mm-4 *M. maximus* 4x.

**Figure 3.**
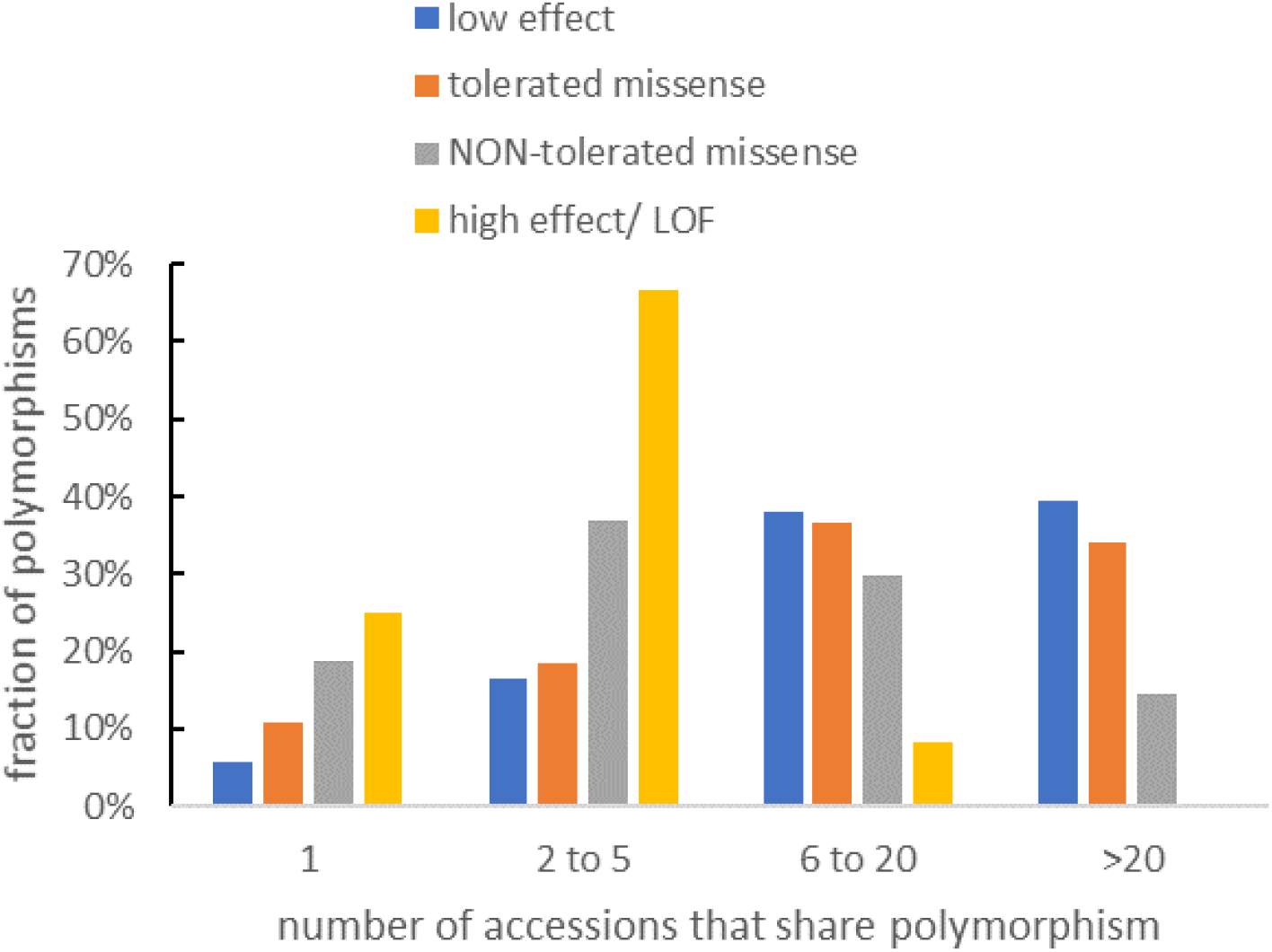
Numbers of accessions that share polymorphisms, grouped by predicted effect.

### Classification of missense variants as tolerated or non-tolerated by SIFT

We downloaded orthologs’ protein sequences for the 20 target genes in angiosperms with fully sequenced genomes from Phytozome v12 (Goodstein *et al*., 2012) and aligned them using Muscle (Edgar, 2004), with default parameters, together with our *U. ruziziensis* reference protein sequence. We assumed that at least one gene must be functional for each of the 50 species, with the exception of BAHD01 and BAHD04 genes, where we included only the 13 commelinid monocot species as their function is believed to be confined to these species (Mitchell *et al*., 2007). We removed any paralogues that did not align well. For each gene, we supplied these alignments and the discovered missense variants to the SIFT web server (Sim *et al*., 2012) . We then used the SIFT prediction to classify the missense variants as tolerated (score >0.05) or non-tolerated (score<=0.05); non-tolerated predictions were all flagged as low confidence because of the small number of sequences available for alignment, while tolerated ones were regarded as reliable.

## Results

### Identification of target genes

We identified genes from the literature, including published work from our own labs, where there was evidence that a loss of function in the gene would confer either increased digestibility (cell wall genes) or increased lipid content in vegetative tissue (lipid genes). This evidence is summarised in Table 1.

We identified orthologues of the genes in Table 1 in *Setaria viridis, Setaria italica, Sorghum bicolor* (as the most closely related fully sequenced genomes) and in the draft genome of *U. ruziziensis* (Supplemental Table 2). We found additional putative paralogues in *U. ruziziensis* for 4CL, CAD, CCoAOMT and CGI58 and we included these in the analysis, naming them with the suffix “_p1”. We therefore had a final total of 11 cell wall genes and 9 lipid genes as our target set identified in *U. ruziziensis*.

RNAseq was carried out on RNA collected from leaves of 104 accessions growing in fields at CIAT. These data have been deposited at NCBI under BioProject PRJNA513453.We took reads from *U. ruziziensis* accessions that mapped to the target *U. ruziziensis* genes (Supplemental Table 2) and unmapped reads and re-assembled these target genes. We compared the resulting sequences with target *U. ruziziensis* and *S. viridis* gene models to improve the *U. ruziziensis* gene models and design baits. We carried out bait capture of genomic DNA from a set of 80 accessions, 74 of which were in the RNAseq set. We used *U. ruziziensis* RNAseq, bait capture DNAseq and Sanger sequencing to iteratively improve the *U. ruziziensis* gene models. We submitted the final gene model versions to Genbank/EMBL and substituted them for the original versions into the *U. ruziziensis* reference genome. We then re-mapped all accessions’ RNAseq and bait capture DNAseq to this updated reference. We called variants on the resulting alignments and identified those that were found in RNAseq and confirmed as present in bait capture DNAseq in same accession or, in cases where DNAseq was not available for same accession, from any other accession of same species. We also looked for special case of loss of splice donor or acceptor in DNAseq, which would be expected to change RNAseq read distribution, but did not find any instances of this. We present results below only for variants found in RNAseq and confirmed in bait capture DNAseq since these have good confidence as the two approaches have different sources of error.

The numbers of variants of different types in the target genes are summarised in Table 2 and the complete set is available in Supplemental Table 3.

We were most interested in mutations that disrupt function but found only 12 variants predicted to lose function (Table 1); however, among the 941 “moderate” mutations (mostly missense nonsynonymous mutations), the single nucleotide polymorphism results in a different amino acid and it is expected that some of these changes will be disruptive. Using the SIFT web server (Sim *et al*., 2012), we supplied our protein alignments of orthologs from fully sequenced plant genomes and used resulting SIFT predictions to categorise missense mutations into tolerated and non-tolerated classes. From this analysis, 256 further variants that we discovered may disrupt gene function (Table 2).

We looked at the number of variants found in individual accessions, grouped by species and ploidy (presented as box whisker plots in Figure 2). As expected, we found more variants in accessions from species that are more distantly related to the *U. ruziziensis* reference, i.e. *U. humidicola* and M. *maximum* (Triviño *et al*., 2017). An outlier accession #26175 for group *U. ruziziensis* with high numbers of variants in these target genes is indicated in Fig. 2 ; this may be misclassified and is probably not *U. ruziziensis* according to a global analysis of all genes’ SNPs (JJDV, unpublished). We found very similar patterns for low effect and tolerated missense polymorphisms suggesting that these both reflect relatedness to the reference. However, we found a different pattern for non-tolerated polymorphisms predicted to affect protein function by SIFT, which were much more common on groups with high ploidy (Figure 2).

Many of the polymorphisms were shared between multiple accessions and we present a summary of this in Figure 3. We found that polymorphisms predicted to disrupt (non-tolerated missense) or eliminate (LOF) function were shared between fewer accessions than other polymorphisms. Polymorphisms characteristic of subgenomes (homeologues) would be expected to be present in all accessions with these subgenomes (typically >20 in the set used here), whereas allelic variants would be present in fewer accessions. From our analysis, it appears that non-tolerated missense and LOF mutations are much more likely to be allelic than homeologous compared with other mutations (Fig. 3).

We manually examined the alignments for the 12 putative LOF alleles we found initially by our automated pipeline (Table 2) and found that 3 were frameshifts in stretches of homopolymer or low complexity, with low coverage in some cases. It is likely that these are real since they were found in the same accessions in RNAseq and gDNA sequencing, but it is also possible they are artefacts due to systematic errors common to both DNA and RNAseq approaches. We found two others were predicted to truncate protein close to C-terminus so were less certain to knock-out function. We therefore designate these 5 as “low confidence” and the remaining 7 as “high confidence”. We show the alignments of these 7 LOF alleles, 3 in cell wall genes (Figure 4), 4 in lipid genes (Figure 5). From a breeding perspective, it is more difficult to transfer an allele from an accession with higher ploidy to a line with lower ploidy; since commercial varieties of these species are tetraploid, this may make the alleles of PXA1, SDP1-like, 4CL_p1 found in accessions with ploidy >4 of less immediate value. This leaves the putative LOF alleles in BAHD01 in tetraploid *U. brizantha*, in CAD_p1 in diploid *U. ruziziensis* and SDP1 in diploid *U. decumbens* as of most potential interest. All these alleles appear to be present in heterozygous form so further breeding would be required even in diploid accessions to achieve complete loss of function.

**Figure 4.**
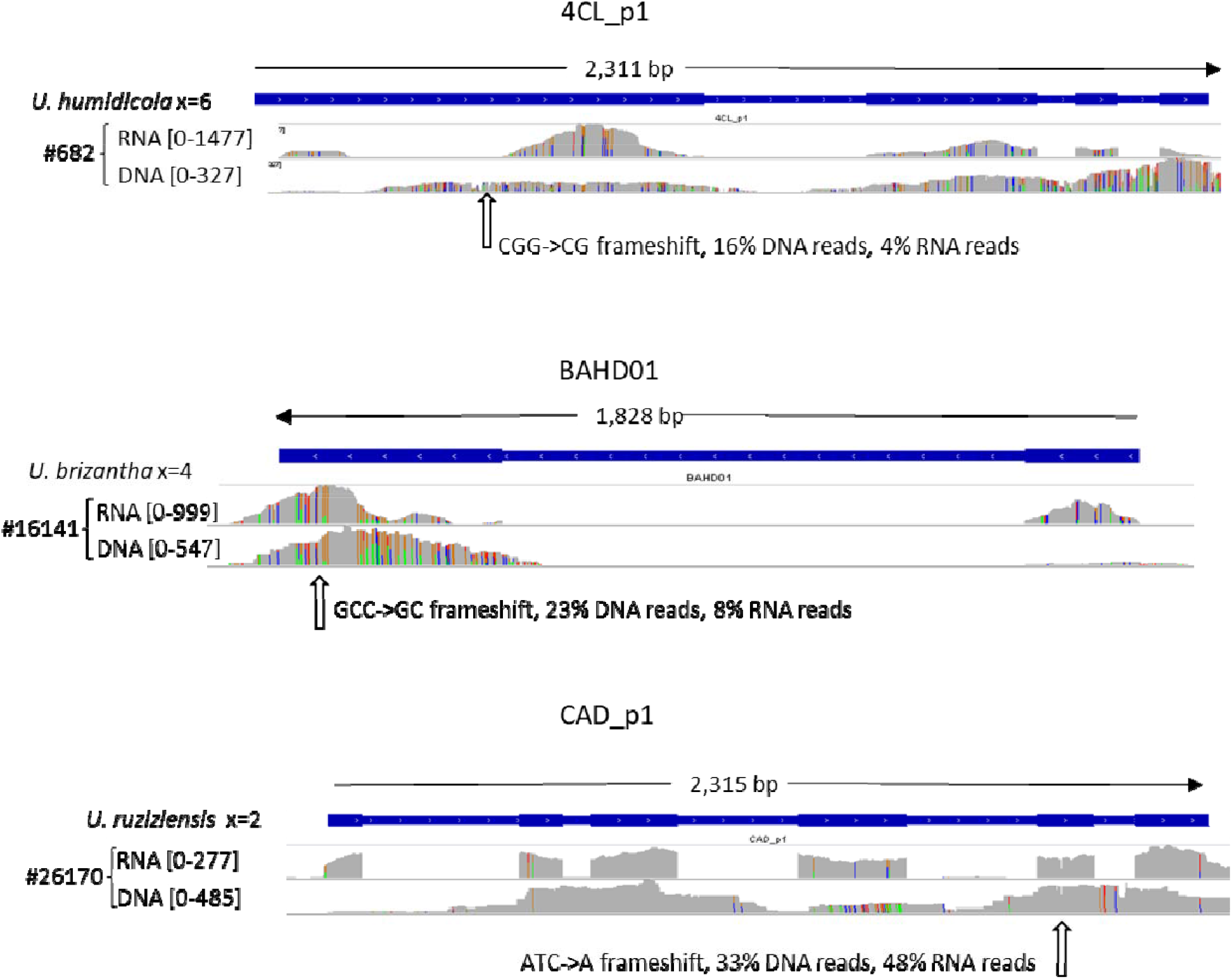
Three cell wall genes with LOF alleles. Read coverage for both RNAseq and DNAseq is shown for accessions indicated which carry LOF allele. Scale of coverage in number of reads and LOF polymorphism are indicated.

**Figure 5.**
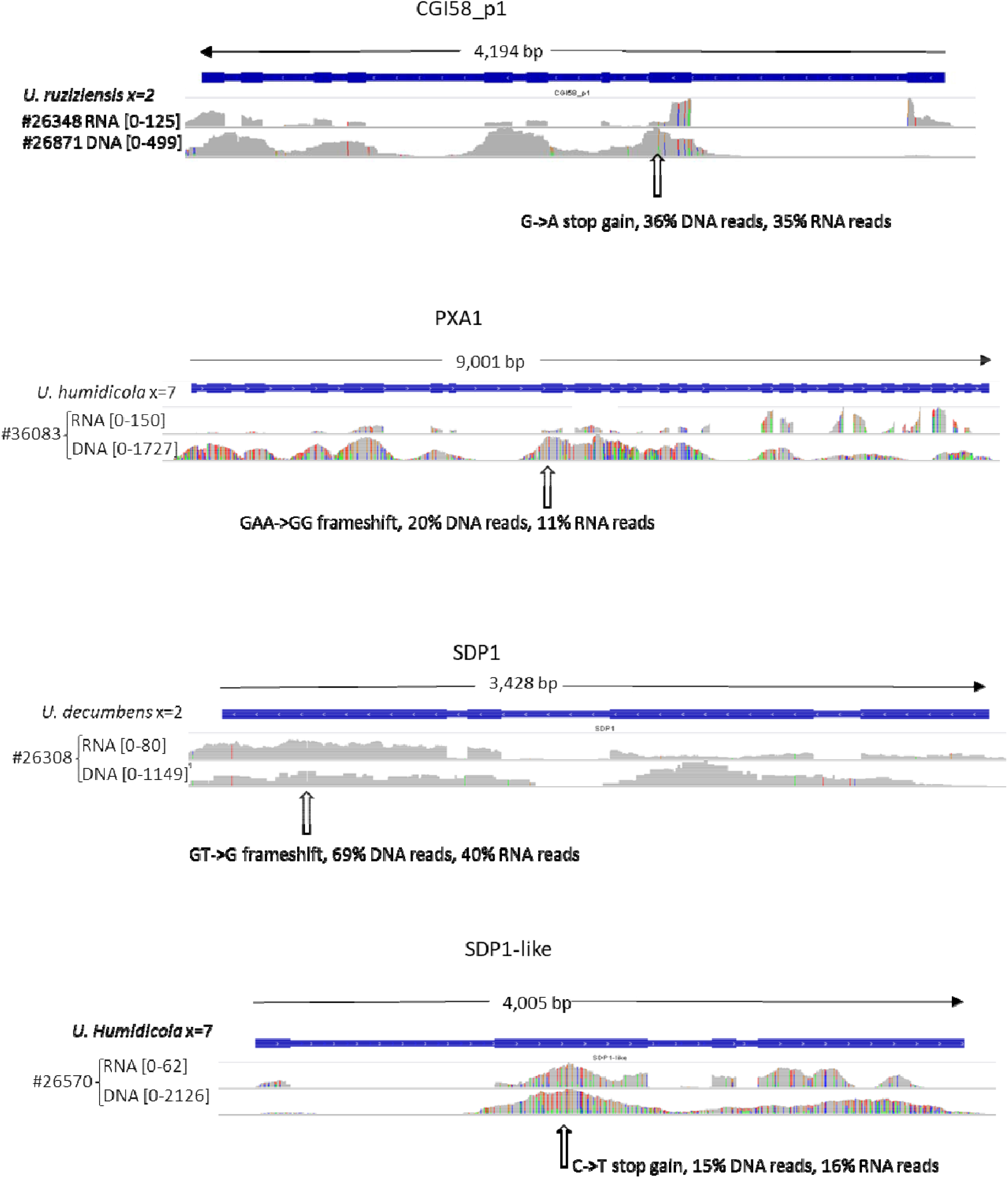
Four lipid genes with LOF alleles. Read coverage for both RNAseq and DNAseq is shown for accessions indicated which carry LOF allele. Scale of coverage in number of reads and LOF polymorphism are indicated.

## Discussion

We developed a new methodology for allele discovery of candidate genes in a collection of diploid and polyploid lines with only a draft genome sequence for one diploid species as reference. Our approach of combining RNA-seq and bait capture (Fig. 1) provides a means of avoiding pseudogenes and resolving complexities. As part of the process, we improved gene models for 18 key genes and confirmed 2 more as accurate in the *U. ruziziensis* v1 genome. Our approach could be adapted for allele discovery in other plant collections or populations.

Our motivation in this work was the hope that breeding tropical forage grass with increased digestibility and lipid content could reduce environmental impact of cattle production by respectively decreasing land requirement and CH_4_ emissions. We selected target genes from our work or the literature where reduction in function improves either digestibility or lipid content of vegetative tissue (Table 1). In the case of digestibility, the evidence comes directly from grass species, whereas the target genes for lipid content have so far only been tested in dicots. These genes differ substantially in the effects of knock-downs or knock-outs and in the evidence for any adverse pleiotropic effects. For some, it is thought that a complete knock-out of function is required for the beneficial effect and this causes little or no side-effects (e.g. COMT in maize; (Vignols *et al*., 1995)). For the BAHD01 and BAHD05 genes putatively involved in addition of hydroxycinnamic acids to arabinoxylan, no complete knock-outs have been reported but knock-downs can have substantial effects (de Souza *et al*., 2018; de Souza *et al*., 2019). In general, LOF alleles have less effect the greater the redundancy from other genes, so are recessive, although dosage effects can occur. In polyploid species like wheat, it can be necessary to stack homozygous LOF alleles in all homeologs to achieve a phenotype (Borrill *et al*., 2019).

We found 941 non-synonymous variants for our 20 target genes within 104 CIAT Genebank accessions confirmed in RNAseq and DNA (Table 2). Most of these likely have little or no effect on function but to gauge which ones are more likely to be detrimental we used the SIFT webserver (Sim *et al*., 2012) to identify a subset of 256 non-tolerated missense variants. Since these are predicted by SIFT based on alignments of all orthologs, they do not reflect relatedness to the *U*. ruziziensis reference and their frequency in accessions was principally dependent on ploidy (Fig. 2). This is most simply explained by the increasing copy number of the genes. An additional effect might be expected where detrimental mutations accumulate in lines with higher ploidy as purifying selection will act less on highly redundant genes, but we could not judge this from our data. In fact, these non-tolerated missense variants were only predicted to be detrimental with low confidence by SIFT due to insufficient diversity of orthologues from fully sequenced plant genomes. These predictions could be improved in future as more genomes are sequenced and knowledge of the proteins improves.

The most secure predictions for disrupted function are the LOF variants with premature stop codons or frameshifts. We found that both non-tolerated missense and LOF variants tended to be shared between fewer accessions compared to other variants (Fig. 3), indicating they were more likely to be allelic than homeologous. Homeologous variants are more ancient than allelic variants so it may be that purifying selection has tended to remove detrimental variants over longer timescales.

On manual inspection of LOF variants, 5 were such that we were not completely confident they were real or were likely to knock-out function. Of the other 7 (Figs. 4, 5), three were of particular interest since they occur in tetraploid or diploid accessions that are more easily incorporated into breeding programmes; these occurred in CAD_p1, BAHD01 and SDP1 genes. However, CAD_p1 is putatively redundant with CAD and we have not found a report of effect of repressing CAD_p1 without also repressing CAD. No complete knock out of BAHD01 has been reported but partial suppression had a substantial effect on digestibility in Setaria (de Souza *et al*., 2018), so it is possible that dosage effects of this allele we found in CIAT accession #16141 might be observed even in tetraploid lines retaining some functional BAHD01 alleles. The SDP1 LOF allele occurs in a diploid *U. decumbens* accession (CIAT #26308) in heterozygous form. This accession is sexual so could be crossed to compatible diploid lines the descendants of which could be crossed to produce a homozygous diploid line. Knock-out of SDP1 alone increases storage lipid content by many fold in vegetative tissue of Arabidopsis, (i.e., it is not redundant with SDP1-like) (Kelly *et al*., 2013), so such a line could be used to test for this effect in *Urochloa* genus. If successful, the line could be crossed into tetraploid commercial breeding populations, e.g. using a chromosome doubling step.

In future, the allele mining approach we describe here could be applied to other genes and need not be confined to alleles detrimental to molecular function. For example, candidate genes underlying apomixis (Worthington *et al*., 2016) and spittlebug resistance (Ferreira *et al*., 2019) traits have recently been identified in *Urochloa*; with improving ability to predict consequences of variants in these, an allele mining approach could be of value. Also, for gene targets such as these where dominant alleles may affect phenotype, candidate gene association genetics could be a useful approach, as successfully applied for the FT gene and flowering time in *Lolium perenne* (Skøt *et al*., 2011). As knowledge of genes improves, allele mining of diverse germplasm will become an increasingly powerful tool to identify lines that could be beneficially brought into many crop breeding programmes.

## Author Contributions

RACM, JJDV and JSHH conceived the project. TKP collected samples with assistance from VC and JA and carried out RNA isolation. SJH carried out DNA sequencing and bait capture. SJH and RACM carried out bioinformatic steps. RACM wrote the manuscript with contributions from SJH, TKP, JJDV, JA, VC, PJE and JSHH.

## Acknowledgements

This work was supported under the RCUK-CIAT Newton-Caldas Initiative “Exploiting biodiversity in Brachiaria and Panicum tropical forage grasses using genetics to improve livelihoods and sustainability”, with funding from UK’s Official Development Assistance Newton Fund awarded by UK Biotechnology and Biological Sciences Research Council (BB/R022828/1). Additional funding for this study was received from the CGIAR Research Programs on Livestock; and Climate Change, Agriculture and Food Security (CCAFS).

## References

Awai K, Xu C, Tamot B, Benning C. 2006. A phosphatidic acid-binding protein of the chloroplast inner envelope membrane involved in lipid trafficking. 103, 10817–10822.

Borrill P, Harrington SA, Uauy C. 2019. Applying the latest advances in genomics and phenomics for trait discovery in polyploid wheat. The Plant Journal 97, 56–72.

Bouvier d’Yvoire M, Bouchabke-Coussa O, Voorend W, Antelme S, Cézard L, Legée F, Lebris P, Legay S, Whitehead C, McQueen-Mason SJ, Gomez LD, Jouanin L, Lapierre C, Sibout R. 2013. Disrupting the cinnamyl alcohol dehydrogenase 1 gene (BdCAD1) leads to altered lignification and improved saccharification in Brachypodium distachyon. The Plant Journal 73, 496–508.

Brenner EA, Zein I, Chen YS, Andersen JR, Wenzel G, Ouzunova M, Eder J, Darnhofer B, Frei U, Barriere Y, Lubberstedt T. 2010. Polymorphisms in O-methyltransferase genes are associated with stover cell wall digestibility in European maize (Zea mays L.). Bmc Plant Biology 10.

Butardo VM, Anacleto R, Parween S, Samson I, de Guzman K, Alhambra CM, Misra G, Sreenivasulu N. 2017. Systems Genetics Identifies a Novel Regulatory Domain of Amylose Synthesis. Plant Physiology 173, 887–906.

Cingolani P, Patel VM, Coon M, Nguyen T, Land SJ, Ruden DM, Lu X. 2012a. Using Drosophila melanogaster as a Model for Genotoxic Chemical Mutational Studies with a New Program, SnpSift. Frontiers in genetics 3, 35–35.

Cingolani P, Platts A, Wang LL, Coon M, Nguyen T, Wang L, Land SJ, Lu X, Ruden DM. 2012b. A program for annotating and predicting the effects of single nucleotide polymorphisms, SnpEff. Fly 6, 80–92.

Comai L. 2005. The advantages and disadvantages of being polyploid. Nat Rev Genet 6, 836–846.

de Souza WR, Martins PK, Freeman J, Pellny TK, Michaelson LV, Sampaio BL, Vinecky F, Ribeiro AP, da Cunha BADB, Kobayashi AK, de Oliveira PA, Campanha RB, Pacheco TF, Martarello DCI, Marchiosi R, Ferrarese-Filho O, dos Santos WD, Tramontina R, Squina FM, Centeno DC, Gaspar M, Braga MR, Tiné MAS, Ralph J, Mitchell RAC, Molinari HBC. 2018. Suppression of a single BAHD gene in Setaria viridis causes large, stable decreases in cell wall feruloylation and increases biomass digestibility. New Phytologist 218, 81–93.

de Souza WR, Pacheco TF, Duarte KE, Sampaio BL, de Oliveira Molinari PA, Martins PK, Santiago TR, Formighieri EF, Vinecky F, Ribeiro AP, da Cunha BADB, Kobayashi AK, Mitchell RAC, de Sousa Rodrigues Gambetta D, Molinari HBC. 2019. Silencing of a BAHD acyltransferase in sugarcane increases biomass digestibility. Biotechnology for Biofuels 12, 111.

Edgar RC. 2004. MUSCLE: multiple sequence alignment with high accuracy and high throughput. Nucleic Acids Research 32, 1792–1797.

Ferreira RCU, Lara LAdC, Chiari L, Barrios SCL, do Valle CB, Valério JR, Torres FZV, Garcia AAF, de Souza AP. 2019. Genetic Mapping With Allele Dosage Information in Tetraploid Urochloa decumbens (Stapf) R. D. Webster Reveals Insights Into Spittlebug (Notozulia entreriana Berg) Resistance. Frontiers in Plant Science 10.

Fu CX, Mielenz JR, Xiao XR, Ge YX, Hamilton CY, Rodriguez M, Chen F, Foston M, Ragauskas A, Bouton J, Dixon RA, Wang ZY. 2011a. Genetic manipulation of lignin reduces recalcitrance and improves ethanol production from switchgrass. Proceedings of the National Academy of Sciences of the United States of America 108, 3803–3808.

Fu CX, Xiao XR, Xi YJ, Ge YX, Chen F, Bouton J, Dixon RA, Wang ZY. 2011b. Downregulation of Cinnamyl Alcohol Dehydrogenase (CAD) Leads to Improved Saccharification Efficiency in Switchgrass. Bioenergy Research 4, 153–164.

Garrison E, Marth G. (July 01, 2012, 2012.) Haplotype-based variant detection from short-read sequencing. ArXiv e-prints.

Gaviria-Uribe X, Bolivar DM, Rosenstock TS, Molina-Botero IC, Chirinda N, Barahona R, Arango J. 2020. Nutritional Quality, Voluntary Intake and Enteric Methane Emissions of Diets Based on Novel Cayman Grass and Its Associations With Two Leucaena Shrub Legumes. Frontiers in Veterinary Science 7.

Giardine B, Riemer C, Hardison RC, Burhans R, Elnitski L, Shah P, Zhang Y, Blankenberg D, Albert I, Taylor J, Miller W, Kent WJ, Nekrutenko A. 2005. Galaxy: A platform for interactive large-scale genome analysis. Genome Research 15, 1451–1455.

Goodstein DM, Shu SQ, Howson R, Neupane R, Hayes RD, Fazo J, Mitros T, Dirks W, Hellsten U, Putnam N, Rokhsar DS. 2012. Phytozome: a comparative platform for green plant genomics. Nucleic Acids Research 40, D1178–D1186.

Gutierrez RA, MacIntosh GC, Green PJ. 1999. Current perspectives on mRNA stability in plants: multiple levels and mechanisms of control. Trends in Plant Science 4, 429–438.

Halpin C, Holt K, Chojecki J, Oliver D, Chabbert B, Monties B, Edwards K, Barakate A, Foxon GA. 1998. Brown-midrib maize (bm1) - a mutation affecting the cinnamyl alcohol dehydrogenase gene. Plant Journal 14, 545–553.

Hufnagel B, Guimaraes CT, Craft EJ, Shaff JE, Schaffert RE, Kochian LV, Magalhaes JV. 2018. Exploiting sorghum genetic diversity for enhanced aluminum tolerance: Allele mining based on the AltSB locus. Scientific Reports 8, 10094.

James CN, Horn PJ, Case CR, Gidda SK, Zhang D, Mullen RT, Dyer JM, Anderson RGW, Chapman KD. 2010. Disruption of the <em>Arabidopsis</em> CGI-58 homologue produces Chanarin–Dorfman-like lipid droplet accumulation in plants. 107, 17833–17838.

Jung JH, Kannan B, Dermawan H, Moxley GW, Altpeter F. 2016. Precision breeding for RNAi suppression of a major 4-coumarate:coenzyme A ligase gene improves cell wall saccharification from field grown sugarcane. Plant Molecular Biology 92, 505–517.

Kannan B, Jung JH, Moxley GW, Lee SM, Altpeter F. 2018. TALEN-mediated targeted mutagenesis of more than 100 COMT copies/alleles in highly polyploid sugarcane improves saccharification efficiency without compromising biomass yield. Plant Biotechnology Journal 16, 856–866.

Kelly AA, van Erp H, Quettier AL, Shaw E, Menard G, Kurup S, Eastmond PJ. 2013. The sugar-dependent1 lipase limits triacylglycerol accumulation in vegetative tissues of Arabidopsis. Plant Physiol 162, 1282–1289.

Kim D, Langmead B, Salzberg SL. 2015. HISAT: a fast spliced aligner with low memory requirements. Nature Methods 12, 357.

Li H, Durbin R. 2010. Fast and accurate long-read alignment with Burrows-Wheeler transform. Bioinformatics 26, 589–595.

Lu B, Xu C, Awai K, Jones AD, Benning C. 2007. A Small ATPase Protein of Arabidopsis, TGD3, Involved in Chloroplast Lipid Import. 282, 35945–35953.

Mitchell RAC, Dupree P, Shewry PR. 2007. A novel bioinformatics approach identifies candidate genes for the synthesis and feruloylation of arabinoxylan. Plant Physiology 144, 43–53.

Mota TR, Souza WRd, Oliveira DM, Martins PK, Sampaio BL, Vinecky F, Ribeiro AP, Duarte KE, Pacheco TF, Monteiro NdKV, Campanha RB, Marchiosi R, Vieira DS, Kobayashi AK, Molinari PAdO, Ferrarese-Filho O, Mitchell RAC, Molinari HBC, D. dos Santos W. 2020. Suppression of a BAHD acyltransferase decreases p-coumaroyl on arabinoxylan and improves biomass digestibility in the model grass Setaria viridis. The Plant Journal n/a.

Nuñez J, Arevalo A, Karwat H, Egenolf K, Miles J, Chirinda N, Cadisch G, Rasche F, Rao I, Subbarao G, Arango J. 2018. Biological nitrification inhibition activity in a soil-grown biparental population of the forage grass, Brachiaria humidicola. Plant and Soil 426, 401–411.

Park SH, Mei CS, Pauly M, Ong RG, Dale BE, Sabzikar R, Fotoh H, Nguyen T, Sticklen M. 2012. Downregulation of Maize Cinnamoyl-Coenzyme A Reductase via RNA Interference Technology Causes Brown Midrib and Improves Ammonia Fiber Expansion-Pretreated Conversion into Fermentable Sugars for Biofuels. Crop Science 52, 2687–2701.

Pellny TK, Lovegrove A, Freeman J, Tosi P, Love CG, Knox JP, Shewry PR, Mitchell RAC. 2012. Cell walls of developing wheat starchy endosperm: comparison of composition and RNA-seq transcriptome. Plant Physiology 158, 612–627.

Robinson JT, Thorvaldsdóttir H, Winckler W, Guttman M, Lander ES, Getz G, Mesirov JP. 2011. Integrative genomics viewer. Nature Biotechnology 29, 24–26.

Saballos A, Ejeta G, Sanchez E, Kang C, Vermerris W. 2009. A Genomewide Analysis of the Cinnamyl Alcohol Dehydrogenase Family in Sorghum [Sorghum bicolor (L.) Moench] Identifies SbCAD2 as the Brown midrib6 Gene. Genetics 181, 783–795.

Saballos A, Vermerris W, Rivera L, Ejeta G. 2008. Allelic Association, Chemical Characterization and Saccharification Properties of brown midrib Mutants of Sorghum (Sorghum bicolor (L.) Moench). Bioenergy Research 1, 193–204.

Sattler SE, Funnell-Harris DL, Pedersen JF. 2010. Brown midrib mutations and their importance to the utilization of maize, sorghum, and pearl millet lignocellulosic tissues. Plant Science 178, 229–238.

Sim N-L, Kumar P, Hu J, Henikoff S, Schneider G, Ng PC. 2012. SIFT web server: predicting effects of amino acid substitutions on proteins. Nucleic Acids Research 40, W452–W457.

Skøt L, Sanderson R, Thomas A, Skøt K, Thorogood D, Latypova G, Asp T, Armstead I. 2011. Allelic Variation in the Perennial Ryegrass <em>FLOWERING LOCUS T</em> Gene Is Associated with Changes in Flowering Time across a Range of Populations. Plant Physiology 155, 1013–1022.

Slocombe SP, Cornah J, Pinfield-Wells H, Soady K, Zhang Q, Gilday A, Dyer JM, Graham IA. 2009. Oil accumulation in leaves directed by modification of fatty acid breakdown and lipid synthesis pathways. Plant Biotechnol J 7, 694–703.

Triviño NJ, Perez JG, Recio ME, Ebina M, Yamanaka N, Tsuruta S-i, Ishitani M, Worthington M. 2017. Genetic Diversity and Population Structure of Brachiaria Species and Breeding Populations. Crop Science 57, 2633–2644.

Vignols F, Rigau J, Torres MA, Capellades M, Puigdomenech P. 1995. The Brown Midrib3 (Bm3) Mutation in Maize Occurs in the Gene Encoding Caffeic Acid O-Methyltransferase. Plant Cell 7, 407– 416.

Villegas D, Arevalo A, Nunez J, Mazabel J, Subbarao G, Rao I, De Vega J, Arango J. 2020. Biological Nitrification Inhibition (BNI): Phenotyping of a Core Germplasm Collection of the Tropical Forage Grass Megathyrsus maximus Under Greenhouse Conditions. Front Plant Sci 11, 820.

Whitehead C, Garrido FJO, Reymond M, Simister R, Distelfeld A, Atienza SG, Piston F, Gomez LD, McQueen-Mason SJ. 2018. A glycosyl transferase family 43 protein involved in xylan biosynthesis is associated with straw digestibility in Brachypodium distachyon. New Phytologist 218, 974–985.

Wijekoon C, Singer SD, Weselake RJ, Petrie JR, Chen G, Singh S, Eastmond PJ, Acharya SN. Down-regulation of key genes involved in carbon metabolism in Medicago truncatula results in increased lipid accumulation in vegetative tissue. Crop Science n/a.

Winichayakul S, Beechey-Gradwell Z, Muetzel S, Molano G, Crowther T, Lewis S, Xue H, Burke J, Bryan G, Roberts NJ. 2020. In vitro gas production and rumen fermentation profile of fresh and ensiled genetically modified high-metabolizable energy ryegrass. Journal of Dairy Science 103, 2405– 2418.

Worthington M, Heffelfinger C, Bernal D, Quintero C, Zapata YP, Perez JG, De Vega J, Miles J, Dellaporta S, Tohme J. 2016. A Parthenogenesis Gene Candidate and Evidence for Segmental Allopolyploidy in Apomictic <em>Brachiaria decumbens</em>. Genetics, genetics.116.190314.

Worthington M, Perez JG, Mussurova S, Silva-Cordoba A, Castiblanco V, Cardoso Arango JA, Jones C, Fernandez-Fuentes N, Skot L, Dyer S, Tohme J, Palma FD, Arango J, Armstead I, De Vega JJ. 2020. A new genome allows the identification of genes associated with natural variation in aluminium tolerance in Brachiaria grasses. Journal of Experimental Botany.

Worthington ML, Miles JW. 2014. Reciprocal Full-sib Recurrent Selection and Tools for Accelerating Genetic Gain in Apomictic Brachiaria. In: Budak H, Spangenberg G, eds. Molecular Breeding of Forage and Turf. Switzerland: Springer International Publishing, 19–30.

Xu C, Fan J, Froehlich JE, Awai K, Benning C. 2005. Mutation of the TGD1 Chloroplast Envelope Protein Affects Phosphatidate Metabolism in <em>Arabidopsis</em>. 17, 3094–3110.

